# Structural basis for curvature generation and functional specialization in spirochete flagella

**DOI:** 10.64898/2026.02.04.703737

**Authors:** Luca Troman, Bindusmita Paul, Jack Kim, J Christopher Fenno, M. Paula Goetting-Minesky, Eric C. Reynolds, Jillian F. Banfield, Paul D. Veith, Debnath Ghosal

**Author notes:** Corresponding author: Debnath Ghosal. These authors contributed equally to this work (Luca Troman, Bindusmita Paul).

## Abstract

Spirochetes are a distinctive phylum of spiral-shaped bacteria, defined by their curved periplasmic flagella, which drive motility by deforming the cell body and enabling efficient corkscrew-like propulsion through viscous environments. Despite their clinical importance, the molecular mechanisms underlying flagellar assembly, curvature and thus motility, remain poorly understood. Here we used cryo-electron microscopy combined with visual proteomic analysis to determine near-atomic resolution structures of two distinct flagellar filaments natively isolated from *T. denticola*, a major oral pathogen. Our structures reveal that filament curvature is generated by the asymmetric decoration of a conserved FlaB core by multiple sheath proteins, including FlaA1/2/3 and two previously uncharacterized proteins, termed FlaL1 and FlaL2. We show that the sheath imposes differential axial compaction on the FlaB core: FlaA1 expands the lattice at the outer curvature, FlaA2/3 compress the lattice at the inner curvature, and FlaL proteins stabilize these asymmetric interactions. Incorporation of distinct FlaB homologs contributes to assembled filament identity, with FlaB3 forming thin filaments and FlaB1/2 interacting with the sheath to form thick filaments. Comparative analysis reveals that FlaL proteins are conserved amongst some *Treponema* species and several other spirochetes, indicating that asymmetric assembly represents a modular solution to the mechanical demands of periplasmic flagella. These findings provide a structural framework for understanding functional specialization in bacterial filaments.

## Introduction

Spirochetes - derived from the Latin *spira* (coil) and the Greek *chaite* (long hair) - constitute a diverse phylum of spiral-shaped diderm bacterium that includes several important human and animal pathogens, such as *Borrelia burgdorferi* (Lyme disease), *Leptospira interrogans* (leptospirosis), *Treponema pallidum* (syphilis) and *Treponema denticola,* a key pathogen in periodontal disease. Across the phylum, the distinctive corkscrew-like motility is central to their pathogenicity, enabling efficient movement through highly viscous environments such as those found in connective tissue^1,2^.

As in many Gram-negative bacteria, motility in spirochetes is driven by flagella. Flagella share a conserved architectural framework comprising a basal body–motor complex, a hook, a filament, and a distal cap. Typically, flagella are assembled through the flagellum-specific type III secretion system which mediates the translocation of flagellin subunits through the central lumen of the growing filament for incorporation at its distal end^3^. Across bacteria, flagellins possess a conserved helical core, while their outer domains are structurally and functionally diverse^4^.

Unlike most bacteria, spirochetes are defined by the unique periplasmic localisation of their flagella, which reside between the outer membrane and the cell body. Although periplasmic and extracellular flagella share common core components, force generation in spirochetes is considerably more complex. Motility arises from the coordinated rotation of bundles of periplasmic flagella anchored at opposite cell poles, which deform the cell body to generate propulsion^1,2^. Despite conservation of core principles, the number, length, and composition of periplasmic flagella vary substantially among spirochete species.

In *T. denticola*, a well-studied model for spirochete motility, propulsion is typically driven by 2-3 periplasmic flagella originating from opposite poles of the cell^5^. These flagella form left-handed helices that wrap around the cell body, which itself adopts a right-handed helical shape^6^. The two flagellar bundles overlap near mid cell, and coordinated rotation of motors with opposing directionality produces undulation of the cell body and the characteristic wave-like swimming motion^1,7^. Biophysical models of spirochete motility suggest that propulsion requires continuous deformation of both the flagella and the cell body, such that elastic forces generated during flagellar rotation are balanced by reactive forces and torques acting on the cell body, resulting in traveling waves along its length^8–10^. These mechanical demands far exceed those placed on extracellular flagella, which do not need to generate force against a deformable cell body.

Consistent with these functional requirements, periplasmic flagella in spirochetes are among the most complex bacterial filaments described. Unlike extracellular flagella, filaments are assemblies of multiple structural components, typically including several flagellin (FlaB) homologues that form the inner core filament, surrounded by additional proteins that assemble into an external sheath^1^. Recent studies further suggest that this sheath may be asymmetrically organized in several spirochete species, potentially contributing to filament curvature and providing a structural basis for motility mechanics^3,11,12^. Nevertheless, the complete molecular organization of periplasmic flagella, and the mechanisms by which curvature is generated and maintained, remain poorly resolved.

To address these outstanding questions, we focus here on the periplasmic flagella of *T. denticola*. *T. denticola* is a highly motile oral spirochete and a major contributor to periodontal disease, one of the most prevalent bacterial infections worldwide and a leading cause of tooth loss. In *T. denticola*, the inner flagellar core is thought to be composed of one or more FlaB homologues—FlaB1 (TDE1477), FlaB2 (TDE1004), and FlaB3 (TDE1475) ^13^. Previous studies have suggested partial functional interchangeability among these FlaB proteins with respect to filament assembly and motility^7^. The outer sheath has been proposed to include FlaA, encoded by TDE1712^14^; however, two additional putative FlaA-like proteins (TDE1408 and TDE1409) have also been identified, though their roles in flagellar assembly and function remain unclear^15^.

Here, we use single-particle cryo–electron microscopy (SPA cryo-EM) of native flagellar filaments from planktonic *T. denticola* cells to determine high-resolution structures of two distinct filament conformations at sub–3 Å resolution. These structures resolve the full lattice organization of the *T. denticola* periplasmic flagellum and provide mechanistic insight into the structural basis of filament curvature and spirochete motility.

## Results

### Planktonic *T. denticola* exhibits two flagella morphologies

To resolve the full structural complexity of flagellar filaments from *T. denticola*, we used cryo-EM single particle analysis (SPA) of native flagella isolated from planktonic cells (Figures 1A-C, S1 and S2). As has been previously observed^7^, our cryo-EM grids displayed two types of flagellar filaments: “thick” containing the FlaA sheath and “thin” comprising solely of the FlaB/flagellin core. We yielded sub-3 Å reconstructions of both flagellar filaments with clearly resolved side chain densities (Figures 1B-C, S2-S6). Both filament types exhibited a conserved 11-protofilament helical arrangement of FlaB subunits repeating every 52 Å, surrounding a 25 Å central lumen (Figures 1B-C).

**Figure 1.**
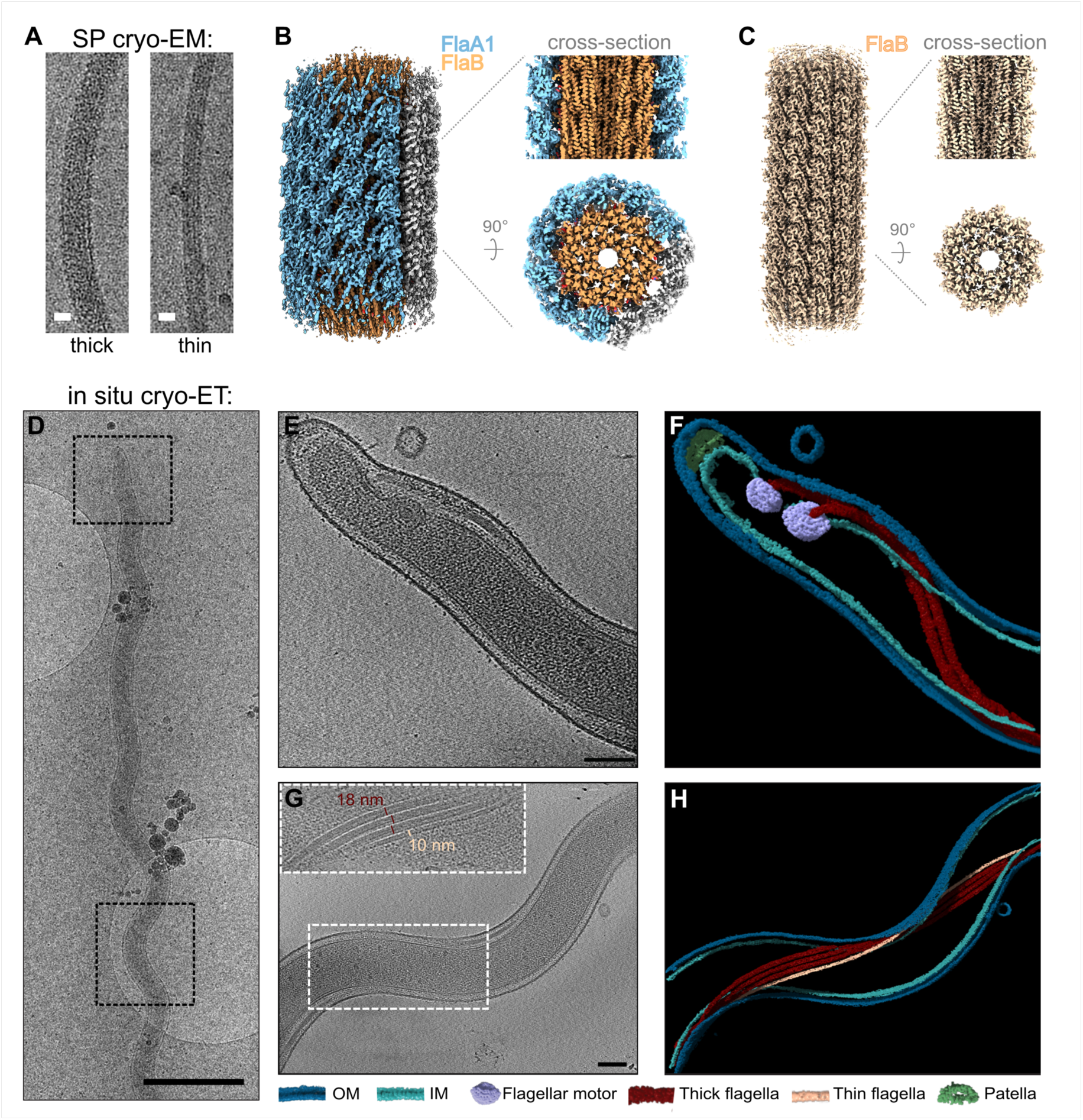
Flagellar filaments in planktonic *T. denticola* cells. (A) Micrographs from single-particle cryo-EM data acquisition showing thick and thin flagellar filaments. (B-C) Sub-3 Å cryo-EM reconstructions for thick (B) and thin (C) filaments. Density colored by known sheath components: FlaB- light orange; FlaA1- light blue; unassigned density- grey. (D) TEM image of a *T. denticola* cell. (E) Cryo-ET of dotted area in (D, top) showing *T. denticola* cell pole. (F) Segmentation of the 3D volume of (E), highlighting the presence of two thick flagellar filaments in the periplasmic space originating from the motors (embedded within the IM) at the cell pole. (G) Cryo-ET of dotted area in (D, bottom) showing mid region of a *T. denticola* cell. Inset, showing the presence of two types of flagellar filament, thick (∼18 nm) and thin (∼10 nm). (H) Segmentation of the 3D volume of (G), highlighting the thick (red) and thin (orange) filaments. Scale bar (A): 10 nm; (D): 1 µm; (E, G): 100 nm.

Although both periplasmic flagella morphologies have been reported in *T. pallidum*^16^ their presence and relevance in intact *T. denticola* remained unclear. To address this, we performed *in situ* cryo-electron tomography (cryo-ET) of planktonic *T. denticola* (Figure 1D-H). Tomograms revealed that most cells possess two periplasmic flagella per pole, each anchored to the inner membrane by a flagellar motor and extending along the length of the cell body (Figure 1E-F). At the midcell region we observed multiple overlapping flagella with two distinct filament diameters: a predominant thick filament (∼18 nm) and a thinner filament (∼10 nm), consistent with the dimensions measured in isolated preparations (Figure 1A, G-H). These data confirm that the two filament morphologies observed by SPA correspond to bona fide periplasmic flagella in intact planktonic cells. We therefore used our high-resolution cryo-EM reconstructions to interrogate the molecular basis of thick filament architecture, with particular emphasis on the composition and organization of the outer sheath.

### Cryo-EM of thick and thin flagellar filaments revealed unique sheath proteins and structural complexity

Intriguingly, the higher resolution reconstruction of the thick filament revealed distinct densities within the outer sheath along the inner curvature (concave side) of the filament as compared to the outer curvature (convex side) (Figures 1B and 2A, shown in grey). This asymmetric distribution is inconsistent with the previously held view that FlaA1 alone constitutes the outer sheath^13^. The sub-3 Å reconstruction enabled direct identification of sheath proteins from the cryo-EM density through the automated model building software ModelAngelo^17^, allowing construction of a full atomic model of the thick filament (Figures 2B, C, S3 and S4). Our near-atomic resolution structure revealed that filament core is composed of FlaB while the outer sheath is dominated by FlaA proteins. In addition to FlaA1 (TDE1712), two FlaA homologues, FlaA2 (TDE1409) and FlaA3 (TDE1408), were identified as integral parts of the flagellar outer sheath (Figure 2B). However, FlaA1/2/3 did not fully account for the observed sheath density (Figure 2B). We also identified two previously uncharacterized proteins that we designate as flagellar lattice proteins (FlaL); FlaL1 (encoded by TDE2349) and FlaL2 (encoded by TDE1480). The identification of unique flagellar sheath proteins was further validated by mass spectrometry of the purified flagellar filaments (Figure 2D, Table S1). For each 52 Å repeat, the outer sheath comprises eight FlaA1, two FlaA2, two FlaA3, and one each of FlaL1 and FlaL2 (Figure 2B). The novel FlaL and putative FlaA proteins form an asymmetric network at the inner curvature of the thick filament (Figure 2C, Movie 1) suggested a direct role in defining filament curvature, which we examine in detail below.

**Figure 2.**
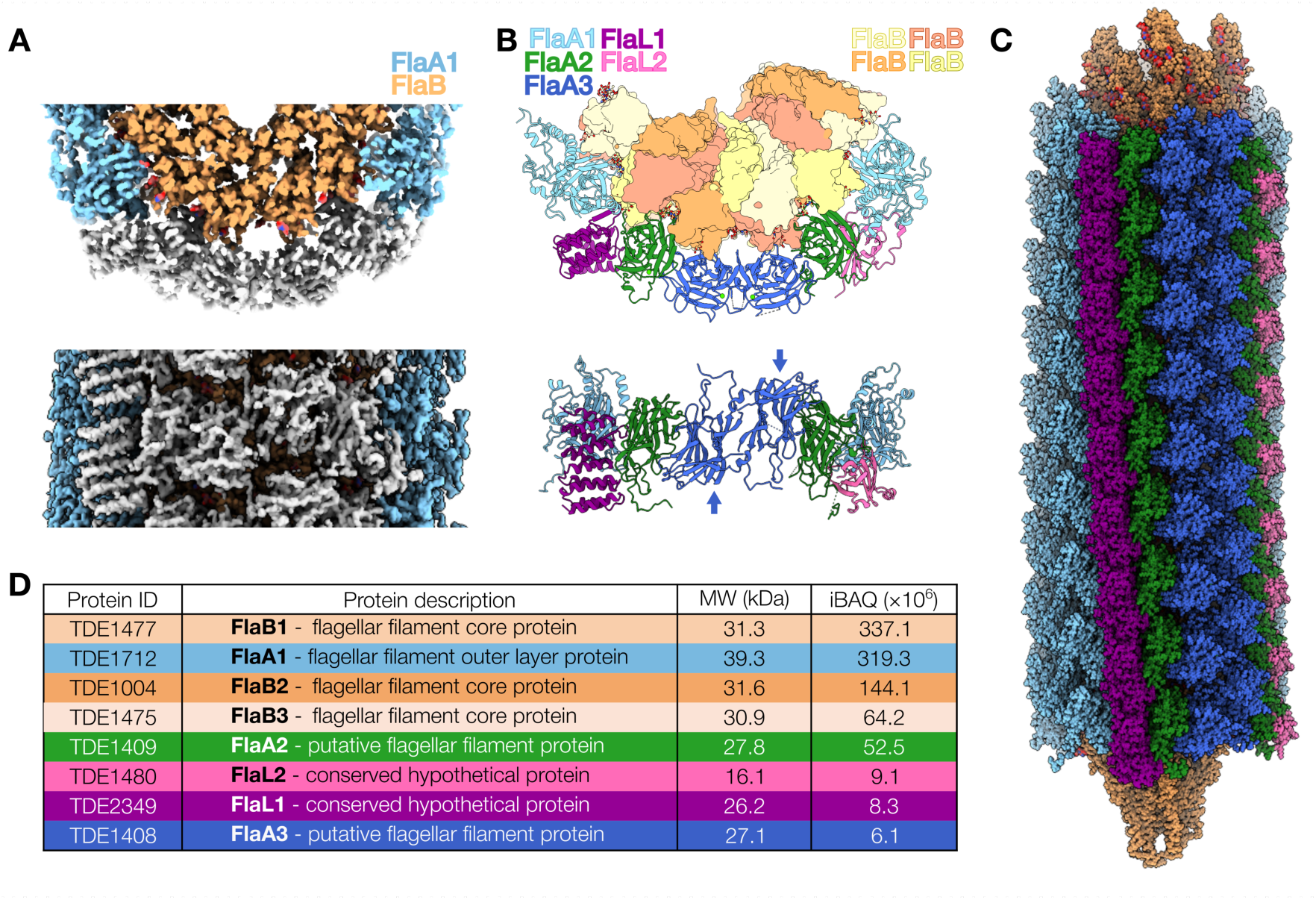
Cryo-EM reveals a complex asymmetric sheath with novel flagellar-lattice proteins. (A) Top-down slice through and side views of the thick filament with unassigned density within the flagellar sheath (grey), FlaA1 (light blue) and FlaB (orange). (B) Fully assigned model of the thick filament containing: FlaB- light yellow, salmon, orange; FlaA1- light blue; FlaA2- green; FlaA3- dark blue; FlaL1- dark magenta; and FlaL2- pink. Arrows indicate the antiparallel arrangement of FlaA3 within the lattice. (C) The full expanded model for the thick periplasmic flagellum containing all components, colored as in (B). (D) The eight most abundant proteins identified by mass spectrometry of the purified flagellar filament sample with absolute abundance of each protein estimated by the iBAQ value.

### Structural specialization of flagellar filaments is driven by distinct FlaB homologues

Our high-resolution SPA reconstructions enabled us to resolve distinct FlaB homologues within the cores of thick and thin flagellar filaments (Figures 3A-B). These represent a second axis of filament specialization that likely cooperates with the sheath to define curvature. Sequence alignments revealed that FlaB homologues - FlaB1, FlaB2, and FlaB3 - share high sequence identity (>70%, Figure S5A), with the most notable sequence differences located at the interface with the outer sheath (Figure 3C). Specifically, FlaB3 features a three-residue longer H4-H5 loop and a four-residue shorter H3-S1 loop compared to FlaB1 and FlaB2 (Figure S5B). These differences enabled automated model assignment of FlaB3 to the thin filament and FlaB2 to the thick filament. These assignments were further validated by manual inspection of the relevant cryo-EM density maps (Figure 3C).

**Figure 3.**
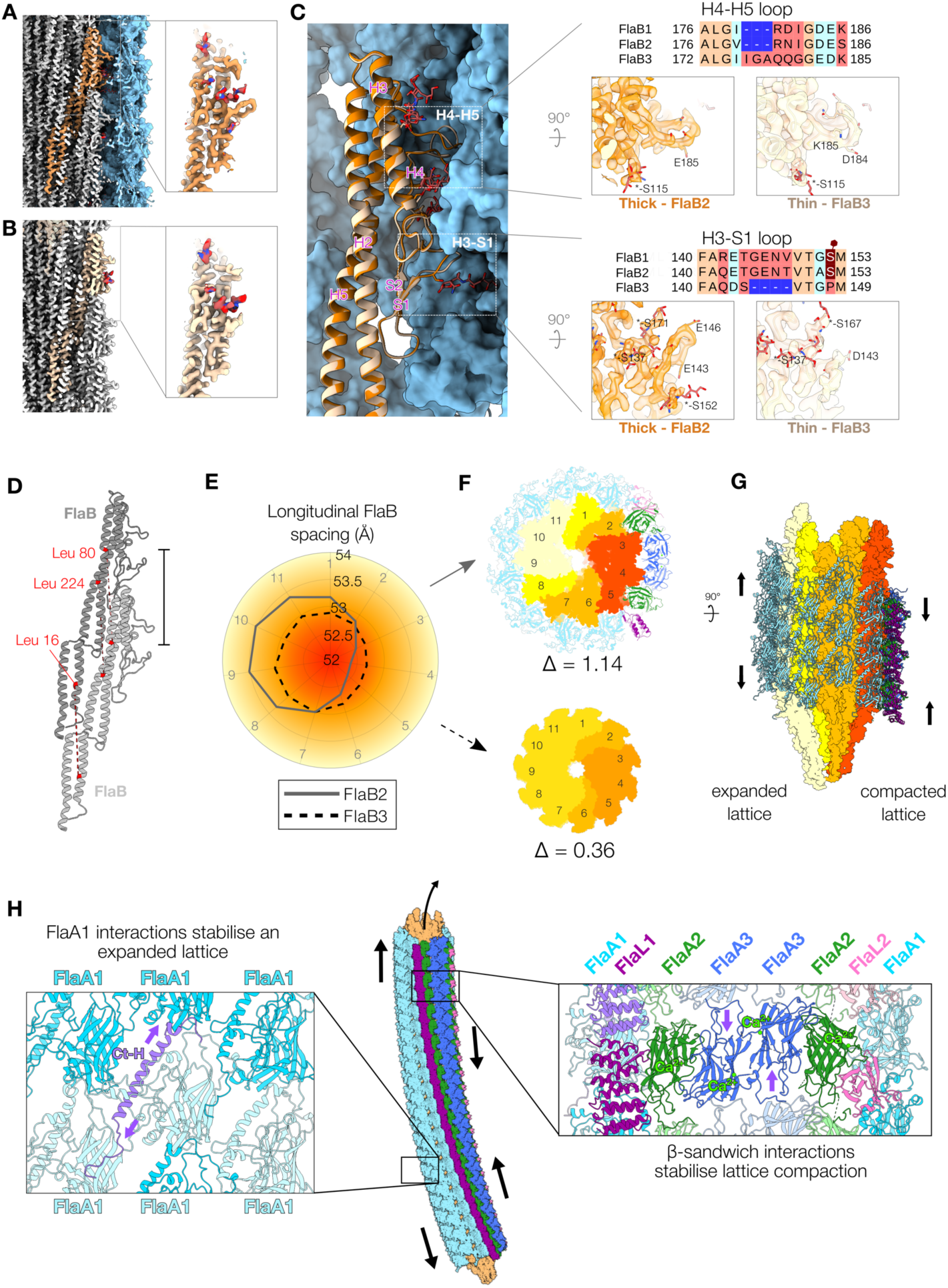
Structural basis of curvature generation in flagellar filament. **(A-B)** Cut through of our cryo-EM reconstructions of the thick (A) and thin (B) flagellar filaments depicting a single FlaB both alone and within the full lattice (FlaB- orange; FlaA1- light blue). (C- left) The molecular interaction between a single FlaB2 (orange) and FlaA1 sheath (light blue) from the thick filament overlaid with the model for FlaB3 in the thin filament (cream). (C- right) Sequence comparison for the H4-H5 loop and H3-S1 loop across all three FlaB homologues with the corresponding FlaB models and cryo-EM density for FlaB from each flagellum. (D) Two longitudinally associated FlaB from our model; marked residues were used for distance measurements along the helical axis. (E) Mean local distance measurements between FlaB monomers within the thick and thin filaments plotted according to position within the lattice. (F) Top-down view of the thick and thin flagella colored according to the measured spacing; higher compaction- red; lower compaction- yellow. (G) The corresponding side view model for the thick flagella. (H) Differential axial compaction mediated through interaction of FlaA1, and FlaL1/FlaA2/FlaA3/FlaL2 with the FlaB core. Diagonal bridging of FlaA1 through the Ct-helix (purple) promotes lattice expansion in the outer sheath whilst at the opposite side of the lattice, associations of tandem intermolecular β-sandwich interfaces in FlaA2-FlaA3-FlaA3-FlaA2-FlaL2 promote lattice compaction.

Previous mass spectrometry studies^18^ have shown that FlaB1 and FlaB2 - but not FlaB3 - are uniquely glycosylated at residues 140-160. Consistent with this report, our cryo-EM density analysis of this region revealed a glycosylated Ser152 in the thick filament, whereas no corresponding density was observed in the thin filament (Figure 3C). While our results confidently annotate that the thin filament is composed primarily of FlaB3 (Figure S6), the thick filament likely contains a heterogeneous mix of FlaB1 and FlaB2. Although our mass spectrometry analysis confirms the presence of both FlaB1 and FlaB2 at high abundance (Figure 2D, Table S1), these two homologues are indistinguishable at our current resolution and are therefore represented as FlaB2 in the thick filament model. Thus, SPA reveals that flagellar filament identity in *T. denticola* is defined by selective incorporation of distinct FlaB homologues: the thin filament exclusively incorporates FlaB3, whereas the thick filament incorporates FlaB1 and/or FlaB2, not FlaB3. These FlaB-specific differences localize predominantly to regions that interface with the outer sheath, consistent with a role for FlaB composition in shaping core–sheath interactions.

### Molecular interaction network in the outer sheath modulates FlaB core structure and defines flagellar curvature

Having established that filament types differ in both core and sheath composition, we analyzed the spatial arrangement of FlaB homologues in each filament type to determine how the molecular interaction network affect filament architecture. FlaB models were fit into non-symmetrized cryo-EM reconstructions, and inter-atomic distances were measured between Cα atoms of three residues across three longitudinally aligned FlaB subunits (Figure 3D). We observed varying degrees of longitudinal lattice compaction, with the thick filament exhibiting approximately threefold greater axial compaction than the thin filament (maximum difference in inter-subunit distance: thick = 1.14 Å; thin = 0.36 Å, Figures 3E-F). Structural mapping of these differences back to our models revealed that in the thick filament, compaction is concentrated within the asymmetric sheath region, whereas the FlaA1-containing sheath region appeared expanded (Figures 3F-G). These findings suggest that the sheath imposes asymmetric mechanical constraints on the FlaB core, generating curvature through differential longitudinal compaction (Figure 3H).

We next analyzed molecular interactions within the completed sheath model (Figure 4, 4A, Movie 2). Previous studies have implicated FlaA1 as a major contributor to flagellar helicity^7^. Our data shows that on the expanded side of the outer sheath, the β-strand-containing N-terminal domain (NTD) of FlaA1 interfaced with three FlaB monomers within the flagellar core (Figure 4B). Each FlaA1 subunit diagonally bridged two neighboring FlaA1s through interaction via the C-terminal α-helix (Ct-helix) (Figure 4C). This Ct-helix extended from the apex of one FlaA1 into a groove formed by the 230–250 loop of an adjacent subunit and continued along the NTD, stabilizing the sheath architecture. These FlaA1-FlaA1 interactions organized and stabilized the outer sheath lattice and transmitted structural cues to the underlying FlaB core, leading to local expansion.

**Figure 4.**
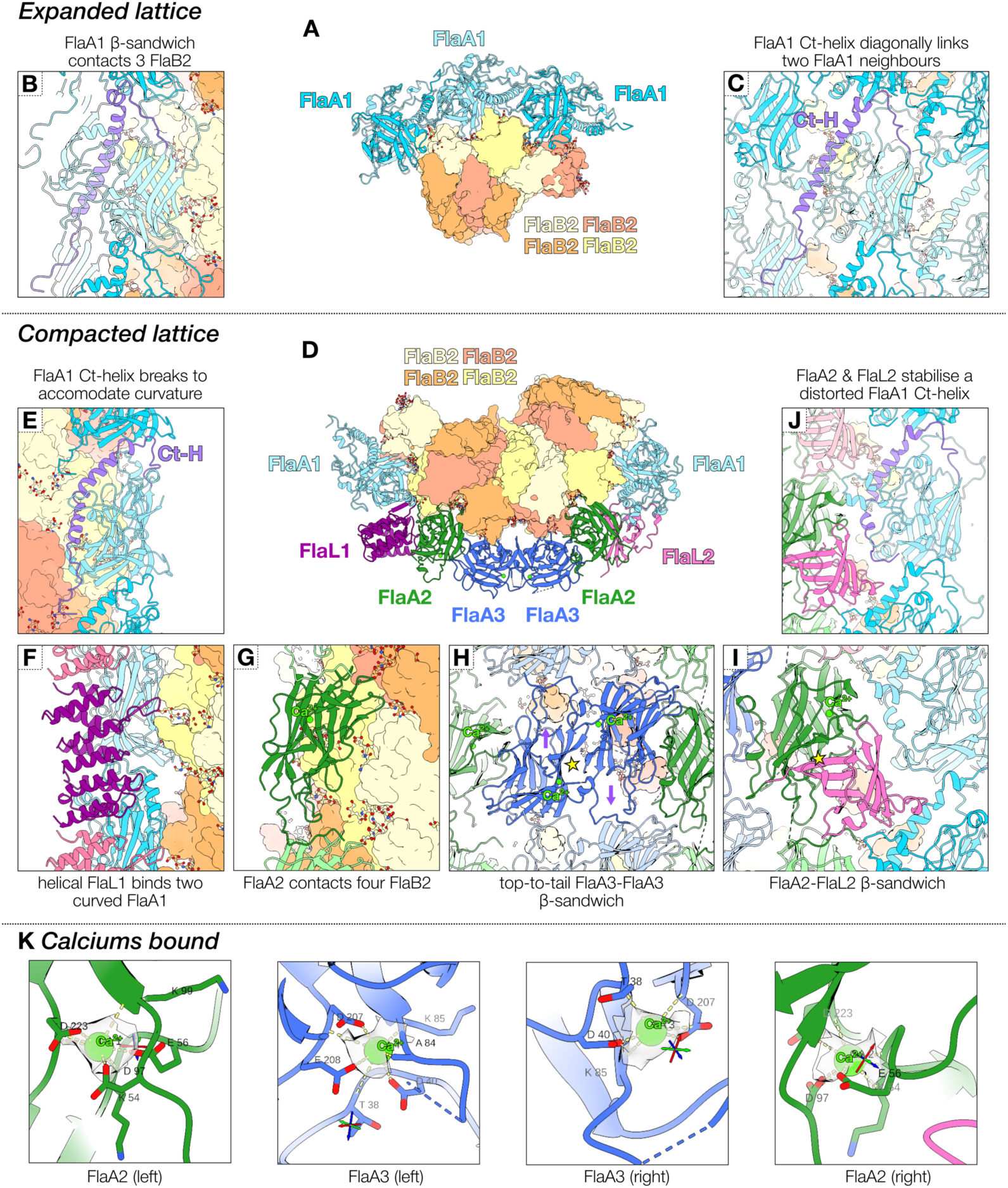
Protein-protein interaction network within thick flagellar filament. (A) Top-down slice through of the outer curvature of the flagellar filament containing FlaA1 within the sheath (blue, cartoon representation) and FlaB within the inner core (surface representation). (B-C) Side views of the filament showing the interaction sites for the FlaA1 with the inner core and neighboring FlaA1 within the sheath. (D) Top-down slice through of the inner curvature of the flagellar filament containing FlaA1, FlaL1, FlaA2, FlaA3, FlaL2 outer sheath components (cartoon representation) and FlaB within the inner core (surface representation). (E-I) Side views of the flagella showing the interaction sites for the outer sheath proteins with the inner core and neighboring proteins within the sheath. (E) The Ct-helix of FlaA1 (purple) distorts to accommodate compaction near the inner curvature of the flagella. (F) FlaL1 (magenta) interacts with two different FlaA1 (light blue) molecules. (G) FlaA2 (green) interacts with four different FlaB at the core. (H) Two antiparallel arranged FlaA3 (dark blue) interact laterally through formation of an intermolecular β-sandwich (yellow star). Longitudinally, the C-termini of each FlaA3 associates with FlaA3 above and below. (I) FlaA2 (green) and FlaL2 (pink) interact through formation of an intermolecular β-sandwich (yellow star). (J) FlaL2 (pink) interacts with FlaA1 via the distorted Ct-helix (purple). (K) Cryo-EM density indicating the presence of Calcium ions (Ca^2+^) within FlaA2 (green) and FlaA3 (dark blue). Each FlaA2/FlaA3 monomer contains one calcium ion per monomer.

In more compacted regions of the sheath (e.g. positions 3–5 in Figure 3F), the FlaA1 Ct-helix was found to be distorted to accommodate lattice compression while preserving FlaA1-FlaA1 contacts (Figure 4E, Movie 2). On the inner curvature of the filament, the underlying FlaB core was more tightly packed, sterically hindering the incorporation of FlaA1 subunits in this region. Consequently, our results indicate that FlaA1 lattice assembly itself induces flagella curvature by locally expanding the FlaB core. This creates spatial asymmetry across the filament, which promotes and stabilizes bending. The two newly identified flagella lattice proteins (FlaL1 and FlaL2) are localized at the interface between the expanded FlaA1 sheath and the compressed FlaA2/3 sheath (Figures 2C, 4D). FlaL1 interacts with the NTD of FlaA1 inducing the ordering of the 230-250 loop into a helical conformation (Movie 2). This structural transition is both spatially and functionally specific to the FlaL1 interaction, suggesting a curvature- or context-dependent recruitment mechanism for selective FlaL1 docking. Notably, with the exception of FlaL1, all other components of this asymmetric lattice are primarily composed of β-sheet assemblies (Figures 4D-I). In contrast, FlaL1 adopts a helical HEAT-repeat domain fold (Figure 4F), a known curvature-sensing motif found in systems such as TOG/Stu2 proteins that recognize curved tubulin conformations during microtubule polymerization^19,20^. FlaL2 binds at the second interface between FlaA2 and the FlaA1 lattice (Figure 4I, Movie 2). It specifically interacts with the distorted C-terminal helix of FlaA1 (Figure 4J, Movie 2), suggesting a role in stabilizing curvature-dependent sheath conformations.

Within the asymmetric sheath region, FlaA2, FlaA3 and FlaL2 interact laterally via tandem intermolecular β-sandwich interfaces (Figures 3H, 4H-I). Both FlaA2 and FlaA3 exhibit density consistent with bound calcium ions, which are known to stabilize β-sandwich folds in bacterial ice-binding adhesins (Figure 4K)^21^. Interestingly, within the lattice, FlaA3 subunits are assembled in a head-to-tail orientation, interacting laterally via inverted β-sandwich interfaces and longitudinally through C-terminal tail anchoring to adjacent FlaA3 subunits (Figure 3H). Both orientations support comparable interfaces with neighboring FlaA2 molecules. FlaA2 is positioned close to the FlaB core, bridging an interface with four FlaB monomers, whereas FlaA3 appears to make only minimal contact with the core, suggesting distinct roles in lattice stabilization and sheath-core integration. Our analysis of the molecular interaction network revealed that the outer sheath induces filament curvature through differential axial compaction of the FlaB inner core (Figure 3H). The key interactions to note are: (i) FlaA1-FlaA1 interactions promote expansion at the outer curvature of the filament, (ii) FlaL1 and FlaL2 bind at the periphery of the expanded FlaA1 lattice and bridge interactions with other sheath components, and (iii) FlaA2 and FlaA3 form a series of calcium-stabilized intermolecular interactions promoting the compression of the FlaB core at the inner curvature of the filament. This balance provides a structural basis for the unique helical motility of this spirochete.

### Conservation of FlaL1 and FlaL2 across the *Spirochaete* phylum

To assess the distribution of newly discovered flagellar lattice proteins (FlaL) across the Spirochaete phylum, we conducted a comprehensive survey of all Spirochaete genomes in NCBI (Figure 5). Our analysis revealed distinct yet partially overlapping patterns of conservation for FlaL1 and FlaL2. FlaL1 is broadly conserved across the *Treponema* genus, exhibiting a markedly broader phylogenetic range than FlaL2, which appears to be less well-conserved within this group. Within *Treponema*, the highest amino-acid identities occur in species closely related to *T. denticola,* including the syphilis-causing *T. pallidum*; outside this core, homologues diverge rapidly.

**Figure 5.**
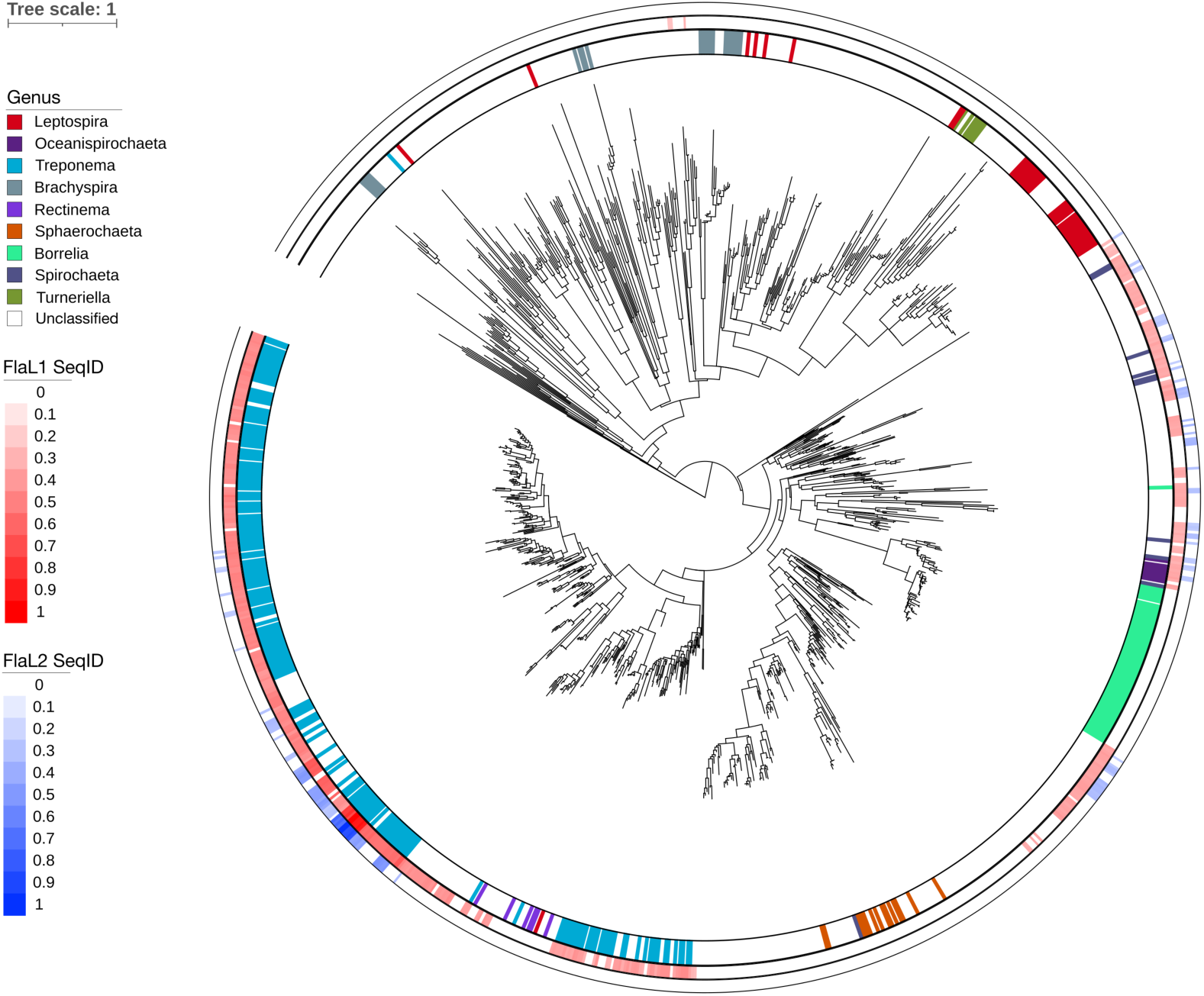
Distribution of FlaL1 and FlaL2 across the *Spirochaete* phylum. A maximum-likelihood tree was built from Rps3 sequences of all publicly available *Spirochaete* genomes (NCBI), with *Elusimicrobium minutum* (GCA_000020145.1) as outgroup. Innermost ring: genus-level taxonomic assignment colored according to the key. Outer rings: amino-acid identity of FlaL1 (red) and FlaL2 (blue) homologues for each genome where a homologue was detected.

Beyond *Treponema,* homologues for both FlaL1 and FlaL2 were identified across the *Spirochaeta*, *Oceanispirochaeta*, and *Borrelia* genera, but none were found in the *Leptospira* genus. While most *Borrelia* genomes lack these genes, a single *B. garinii* genome harbors a FlaL1 homologue, and several unclassified spirochaete genomes branching with *Borrelia* encode both FlaL1 and FlaL2. Notably, the sequence divergence of FlaL1 and FlaL2 homologues does not correlate with the overall species divergence based on the Rps3 phylogeny. This suggests that their conservation is linked to functional importance within these species rather than evolutionary relatedness, as the evolutionary paths of these loci appear to be decoupled from overall species divergence.

## Discussion

Here, using a combination of cryo-EM SPA and proteomic analysis, we resolved the architecture of the periplasmic flagella within the oral pathogen *T. denticola* at unprecedented resolution. Near-atomic resolution structures of the planktonic spirochete flagella revealed striking levels of complexity and established that spirochete flagellar curvature can be encoded by asymmetric decoration of an otherwise conserved flagellin core, rather than by intrinsic curvature of the flagellin filament itself.

### Assembly and curvature generation in *T. denticola* flagella

In both exoflagellates and spirochetes the core flagellar assembly is coordinated by a flagellar basal body or motor, a hook junction and a distal capping complex^22^. Core flagellins (FlaB1/2/3) are secreted through a central lumen and incorporated at the filament tip, assisted by a capping complex (FliD)^23^. Consistent with this model, our cryo-EM reconstructions revealed a central lumen in thick and thin filaments. In contrast, outer sheath components carry N-terminal signal peptides for the Sec-dependent periplasmic secretion indicating that sheath assembly occurs independently of core filament growth. How sheath assembly is coordinated, and whether it is templated by the hook junction (FlgE/FlgK/FlgL), remains unknown.

In motile *T. denticola*, thick flagella display asymmetric compaction of the FlaB core, driven by non-uniform decoration of the outer sheath. We discovered two previously uncharacterized sheath proteins, FlaL1 and FlaL2, bridging the expanded FlaA1-rich outer curvature and the compacted FlaA2/3-rich inner curvature. FlaL1 consists of a HEAT repeat domain, which is known to function as a curvature sensor in other systems - for example, TOG/Stu2 proteins in eukaryotes recognize curved tubulin conformations during microtubule polymerisation^19,20^. FlaL2 binds distorted FlaA1 Ct-helices, stabilizing the compressed FlaA1 conformation that results from the regional lattice compaction. Together, they promote continuous sheath coverage and define filament curvature arising from the sheath-induced deformation of the flagellin core through differential axial compaction of FlaB. The asymmetric lattice we describe provides a structural solution to the unique mechanical challenge faced by periplasmic flagella, which must continuously bend both the filament and the cell body to generate propulsion.

### FlaB as a potential regulatory switch

High resolution cryo-EM SPA revealed that the thick and thin flagella contain distinct homologues of the FlaB flagellin suggesting regulated incorporation rather than a stochastic process. The thin flagellar filaments selectively incorporate FlaB3, whereas thick flagellar filaments incorporate FlaB1/2. This contrasts with a previous report of interchangeability among FlaB homologues^7^. Previous *T. denticola* studies^24^ have linked higher FlaB3 expression with lower levels of motility and a phenotype resembling the protruding flagella observed in older/biofilm cultures^6^. This is consistent with our observations and suggests that FlaB incorporation may serve as a regulatory mechanism modulating filament architecture, curvature, and function. Thus, FlaB specialization is necessary for filament-type identity and sheath compatibility, but asymmetric sheath assembly is sufficient to impose curvature on the FlaB core. How the same flagellar motor selectively incorporates distinct FlaB homologues remains an intriguing question for future investigation.

### Conservation and variation of newly discovered FlaLs among spirochetes

The patchy conservation of FlaL homologues across spirochetes suggests that asymmetric sheath-mediated curvature is a modular solution that can be adapted or replaced by alternative mechanisms in different lineages. While newly discovered FlaL1/2 proteins are less conserved outside of *Treponema species* (Figure 5), similar curvature-inducing principles may apply across other spirochetes. Previous literature has shown that in *Leptospira*, sheath components (FlaA2 and FlaAP) localize at the inner curvature of the flagella^12^, mirroring the roles of *T. denticola* FlaA2/3 and FlaL1/2. Furthermore, *Leptospira* FlaA2 selectively binds glycosylated FlaB homologues, supporting the notion that filament assembly could be controlled at the core-flagellin level with the aid of differential glycosylation.

Together, our results show that spirochete flagellar curvature is established by an asymmetric sheath lattice acting on a conserved FlaB core, rather than arising from intrinsic curvature of the filament itself. This provides a structural solution to the unique mechanical demands of periplasmic flagella, which must continuously bend both the filament and the cell body to generate motility. The identification of FlaL1 and FlaL2 as curvature-associated components highlights a modular mechanism that may be adapted across spirochetes and offers a framework for understanding how filament architecture can be modulated by sheath composition. More generally, these findings illustrate how complex bacterial filaments can achieve functional specialization through combinatorial assembly of conserved subunits and accessory proteins.

## Acknowledgements

This project was supported by an NHMRC grant (APP1196924) and Cumming Global Centre for Pandemic Therapeutics, Peter Doherty Institute for Infection and Immunity grant (CGCPT00060) awarded to DG, NHMRC grant APP1193647 to ECR, Australian Dental Research Foundation grant (0276-2022) awarded to BP, DG, PDV and ECR and US Public Health Service grant DE025225 to JCF. BP is supported by the Melbourne Research Scholarship. We also acknowledge the Ian Holmes Imaging Centre for access to microscopy facilities, and the Melbourne Mass Spectrometry and Proteomics Facility at the Bio21 Institute for access to mass spectrometry instrumentation.

## Authors contributions

Conceptualization, D.G., L.T, B.P.; study design, D.G., L.T., B.P., P.D.V., E.C.R.; methodology, L.T., B.P., J.K., J.C.F., M.P.G-M., J.F.B., P.D.V., D.G.; formal analysis, L.T., B.P., J.K., J.C.F., M.P.G-M., J.F.B., E.C.R., P.D.V., D.G.; investigation, L.T., B.P., J.K., J.C.F., M.P.G-M., J.F.B., E.C.R., P.D.V., D.G.; data curation, L.T., B.P., J.K., J.C.F.; writing – original draft, L.T., B.P., and D.G.; writing – review & editing, all authors; funding acquisition, D.G., E.C.R., J.F.B., J.C.F., P.D.V.

## Data availability

Cryo-EM reconstructions of the outer curvature of the thick flagellar filament, inner curvature of the thick filament, and thin flagellar filament are deposited under EMDB-71672, EMDB-71673, and EMDB-71671, respectively; corresponding atomic models are available in the Protein Data Bank under accession codes PDB 9PIM, 9PIN, and 9PIL. The mass spectrometry proteomics data have been deposited to the ProteomeXchange Consortium via the PRIDE partner repository with the dataset identifier PXD066463.

## Materials and Methods

### Bacterial strains and growth

*Treponema denticola* ATCC 35405 was used in this study. The cultures were grown at 37 °C in a MACS MG500 anaerobic workstation (Don Whitley Scientific, U.K.). *T. denticola* was grown in oral bacteria growth medium (OBGM) as described previously^25^.

### Cryo-electron tomography sample preparation

Planktonic cell suspensions (OD_650_ ∼0.2) were mixed with 10 nm colloidal gold beads (Sigma-Aldrich, Australia) precoated with BSA. The sample was pipetted onto glow discharged copper or gold R2/2 Quantifoil holey carbon grids (Quantifoil Micro Tools GmbH, Jena, Germany). Freezing and blotting were done using a Vitrobot IV (FEI Thermo Fisher Scientific) with chamber humidity set to 100%. Samples were blotted for 4 seconds with a blot force of 8 before being plunge frozen in liquid ethane.

### Cryo-electron tomography data collection and data processing

Grids were imaged using Titan Krios G4 cryo-EM operating at 300 kV acceleration voltage, and a K3 direct detector equipped with a Gatan bio quantum energy filter (slit width 20 eV). Tilt-series were acquired with dose symmetric tilt scheme using Tomography 5 software version 5.14 (Thermofisher Scientific) between tilt range of ±51° with 3° increments. Data were collected with a total fluence of 120 e^-^/Å^2^, a defocus of -8 μm, and pixel size of 3.39 Å/pixel.

Tilt-series alignment was performed using the IMOD software package^26^. Initially, x4 binned tilt-series (13.56 Å/pixel) were aligned using IMOD. The aligned tilt-series were then used for tomogram reconstruction using Tomo3d^27^.

### Tomogram segmentation

For the better visualization of the tomograms, 3D volumes were segmented using Dragonfly software (version 2022.2)^28^. After import into Dragonfly, tomograms were filtered using built-in filters to increase clarity. Followed by manual segmentation of 15 to 20 slices from individual tomograms. These manually segmented slices were used as inputs for neural network-based training using the U-Net architecture with a 2.5D input dimension (5 slices)^29^. After iterative training, the models were used to segment tomograms, with manual corrections implemented where necessary. Built in functions within Dragonfly software were used to generate 2D images and 3D movies.

### Periplasmic flagellar filament purification

500 ml *T. denticola* grown to late-log phase was harvested by centrifugation at 15,000 × g for 30 mins at 4°C. The cell pellet was washed with 10 mM Na phosphate buffer (pH 5.3) and resuspended in the same buffer. To detach the periplasmic filaments, the cell suspension was passed six times through a 25-gauge hypodermic needle to apply shear force. Cell debris were removed by two rounds of centrifugation at 15,000 × g for 30 mins at 4°C. The resulting supernatant was then ultracentrifuged at 150,000 × g for 4 h at 4°C to isolate the crude filament fraction, which was resuspended in 10mM Tris buffer (pH 7.5, 100 mM NaCl). For further purification, the crude fraction was run through velocity gradient centrifugation in a 10-50% OptiPrep gradient at 150,000 × g for 5 h at 4°C. Pulled fractions were analyzed by SDS-PAGE to identify fraction containing periplasmic flagella based on their expected molecular weight profile (Figure S1A). Negative staining and TEM imaging were performed on all fractions to confirm the presence of periplasmic flagella. The identified fraction was then concentrated, and OptiPrep was removed by centrifugation at 150,000 × g for 4 h at 4°C, followed by resuspension in 10 mM Tris buffer (pH 7.5, 100 mM NaCl).

### Single particle cryo-EM sample preparation

Shortly before use Quantifoil R1.2/1.3 holey carbon grids (Quantifoil Micro Tools GmbH, Jena, Germany) were glow-discharged. Subsequently, the grid was loaded into the humidity chamber of the Leica EM GP2 held at 95% humidity and 4°C. 4 µL of purified *T. denticola* filaments were applied to grid and excess liquid was removed by blotting with filter paper from the back of the grid (with the Leica EM GP2 sensor enabled distance of 1 mm and blotting time of 5 sec) before immediate vitrification in liquid ethane.

### Cryo-EM single particle data collection

High-resolution data for single-particle cryo-EM analysis were collected on FEI Titan Krios G4 300 keV FEG transmission electron microscope (Thermo Fisher Scientific). Micrographs were recorded using a K3 Summit direct electron detector (Gatan) equipped with a post-column energy filter set to a slit width of 20 eV. Data were acquired using automated EPU software at a nominal magnification of 105,000 × with pixel size of 0.834 Å and a defocus range of -0.5 to -1.8 µm (Table S2).

### Cryo-EM single particle data processing

With the exception of particle picking, all image processing steps were carried out using RELION-5^30^. Beam-induced motion within each movie was corrected using MotionCorr2^31^, and the parameters of the contrast transfer function (CTF) for each motion-corrected, dose-weighted movie were determined using CTFFIND4^32^, both within the RELION interface. The resulting micrographs were imported into crYOLO (v1.9.9)^33,34^ for automated filament picking. Manually picked filaments from 33 micrographs were used to train an initial model to pick both thick and thin filaments and avoid overlapping areas. The model was used to pick filaments from the entire dataset with intervals of 62 pixels between boxes (∼52 Å), minimum length of 8 particles and an optimized threshold of 0.4.

The resulting coordinates were re-imported into RELION-5 with a box size of 512 × 512 pixels (427 × 427 Å) and 4× binning to 128 × 128 pixels. Initial classification mirrored microtubule processing methods^35,36^. In brief, 4× binned particles were subjected to supervised 3D classification for a single iteration into two classes with references as the tomographic reconstructions of the thick and thin filaments low pass filtered to 20 Å. Particles assigned into each class were separated into “thick” and “thin” for downstream processing.

A 2D classification of the roughly aligned particles (with no image alignments) allowed removal of overlapping filaments and a further 3D classification allowed removal of any particle lattice assemblies. Next, the separated “thick” and “thin” particles were re-extracted with the full box size of 512 pixels (0.833 Å/pixel) for RELION 3D refinement. For the thick filament, a focused 3D classification with masking of the asymmetric region (no image alignments and high T threshold) separated classes according to heterogeneity in this part of the lattice. One class representative of approximately a quarter of the data resolved well, these particles were taken forward for further refinement. For both thick and thin filament reconstructions data was further refined through iterations of CTF refinement and Bayesian Polishing. In the thick filament, resolutions for different regions were further improved independently: (i) the asymmetric lattice was improved through local masking, and (ii) the FlaA1 outer sheath was refined through RELION symmetry expansion (twist -33.57°, rise 23.9 Å, asu 6) combined with focused masking. For the final reconstitution of the thin filament, helical symmetry (11 asu, twist 65°, rise 4.8 Å) was imposed during the RELION refinement. RELION post-processing of unsharpened half-maps from each refinement were used to generate Fourier Shell Correlation (FSC) analysis to describe resolution. Finally, deepEMhancer^37^ of the unsharpened half maps were used for local sharpening to improve the interpretability for model building.

### Protein identification, model generation and model refinement

The final post-processed reconstructions were analyzed using ModelAngelo^17^ to search against the *T. denticola* proteome (Proteome ID #UP000008212). Custom scripting was used to process the output and identify the fragments. ModelAngelo was then rerun using a FASTA file containing the amino acid sequences for FlaA1, FlaA2, FlaA3, FlaB1, FlaB2, FlaB3, FlaL1 and FlaL2 generating initial models for each chain that served as starting models for further refinement. Missing residues were built in Coot^38^, and the model was subsequently refined in ISOLDE^39^ in ChimeraX^40^ using interactive, force field-based molecular dynamics fitting against both the DeepEMhancer post-processed density and the unsharpened maps from RELION 3D auto-refinement. According to previous literature^18^, custom glycans were first drawn in ChemDraw and exported in SMILES format. Restraints were generated using Phenix (version 1.21.2)^41^, and the ligands along with their restraint files were imported into Coot and docked into the density at the appropriate sites. Finally, ligands were covalently linked to the FlaB using phenix.ligand_linking^41^. The full model was then further refined through multiple iterations of Phenix real space refinement, ISOLDE^39^ and Coot^38^. For modelling within Phenix: only unsharpened maps were used during real space refinement, with non-crystallographic symmetry restraints applied across core flagellins and across each set of three longitudinal repeating sheath components within each model.

### Lattice compression measurements

Models for FlaB were individually rigid body docked into the best C1 reconstructions of the full thick or thin filaments within ChimeraX^40^. In total 33 individual models of FlaB were fit into each filament type representing three repeats of 11 asymmetric subunits/protofilaments. Longitudinal spacing was assessed by measuring distances in ChimeraX^40^ between Cα atoms of three reference residues (16, 80, and 216) across two sets of longitudinally adjacent FlaB monomers, yielding six measurements per FlaB protofilament. The mean intermolecular distances within each protofilament were represented within a circle plot generated using MATLAB (version 24.2.0 (R2024b))^42^. FlaB within each model was then colored in ChimeraX^40^ along a gradient reflective of mean spacing measurements within each protofilament.

### Proteomic sample preparation

For protein identification, the purified periplasmic flagellar filament sample was dissolved in 1× NuPAGE LDS sample buffer containing 50 mM dithiothreitol to denature proteins. The sample was then heated and subjected to SDS-PAGE for 6 minutes, allowing it to migrate just beyond the well. The corresponding protein band was excised, and in-gel trypsin digestion was performed as previously described^43^.

Peptides were extracted sequentially: first with 0.1% aqueous trifluoroacetic acid (TFA) and then with 50% acetonitrile–0.1% aqueous TFA, each for 15 minutes in an ultrasonication bath. The combined extracts were evaporated using a vacuum centrifuge and dissolved in 2% acetonitrile–0.1% aqueous TFA for mass spectrometry (MS) analysis.

### LC-MS/MS data collection and analysis

The digests were analyzed by LC-MS/MS using an Orbitrap Eclipse mass spectrometer coupled to an Ultimate 3000 UHPLC system (Thermo Fisher Scientific). Buffer A was 0.1% formic acid (FA), 2% acetonitrile in H_2_O, and buffer B was 0.1% FA in acetonitrile. Sample volumes of 2 µL were loaded onto a PepMap C18 trap column (75 µM ID x 2 cm, 3 µM, 100 Å, Thermo Fisher Scientific) and desalted at a flow rate of 5 µL min^-1^ for 6 min using buffer A. The samples were then separated through a PepMap C18 analytical column (75 µM ID x 15 cm, 2 µM, 100 Å, Thermo Fisher Scientific) at a flow rate of 300 nL min-1, with the percentage of solvent B in the mobile phase changing from 3% to 23% in 29 min, from 23% to 40% in 10 min, and from 40% to 80% in 5 min. The spray voltage was set at 1.9 kV and the temperature of the ion transfer tube was 275°C. The full MS scans were acquired over a m/z range of 375-1500, with a resolving power of 120,000, and an automatic gain control (AGC) target value of 4.0x105 and a maximum injection time of 50 ms. Dynamic exclusion was set at 30 s. Higher-energy collisional dissociation MS/MS scans were acquired within a 3 s cycle time at a resolving power of 15,000, AGC target value of 5.0x104, maximum injection time of 22 ms, isolation window of m/z 1.6 and collision energy of 30%. All spectra were recorded in profile mode.

Raw MS files were analyzed by MaxQuant (v2.4.2.0)^44^ using label-free quantification (LFQ) searching against a *T. denticola* ATCC 35405 protein sequence database^45^ and a “contaminants” database provided by MaxQuant. The default recommended parameters were used for LFQ except iBAQ quantitation was included and oxidation (Met) was the only optional modification used. The false discovery rate was the default setting of 0.01 (1%). The iBAQ metric was used to estimate the absolute abundance of the flagellar-associated proteins.

### Phylogenetic tree

Genomes classified as *Spirochaetota* were downloaded from National Center for Biotechnology Information (NCBI)^46^, dereplicated at 95% average-nucleotide-identity (ANI) with dRep (v3.4.0)^47^ and filtered to ≥ 90% completeness and ≤ 5% contamination using CheckM2 (v1.0.1)^48^. Ribosomal protein S3c (RPS3c) was detected in the high-quality genomes with HMMER (v3.3.2) hmmsearch^49^, the recovered RPS3c sequences were aligned in Clustal Omega (v1.2.4)^50^, and a maximum-likelihood tree was inferred with IQ-TREE 2 (v2.2.0)^51^. FlaL1 and FlaL2 homologues were detected across *Spirochaetota* genomes using MMseqs2^52^. Phylogenetic tree was visualized with Interactive Tree of Life (iTOL v7.2.1)^53^.

