## Supplementary material for "Structural basis for curvature generation and functional specialization in spirochete flagella": SI

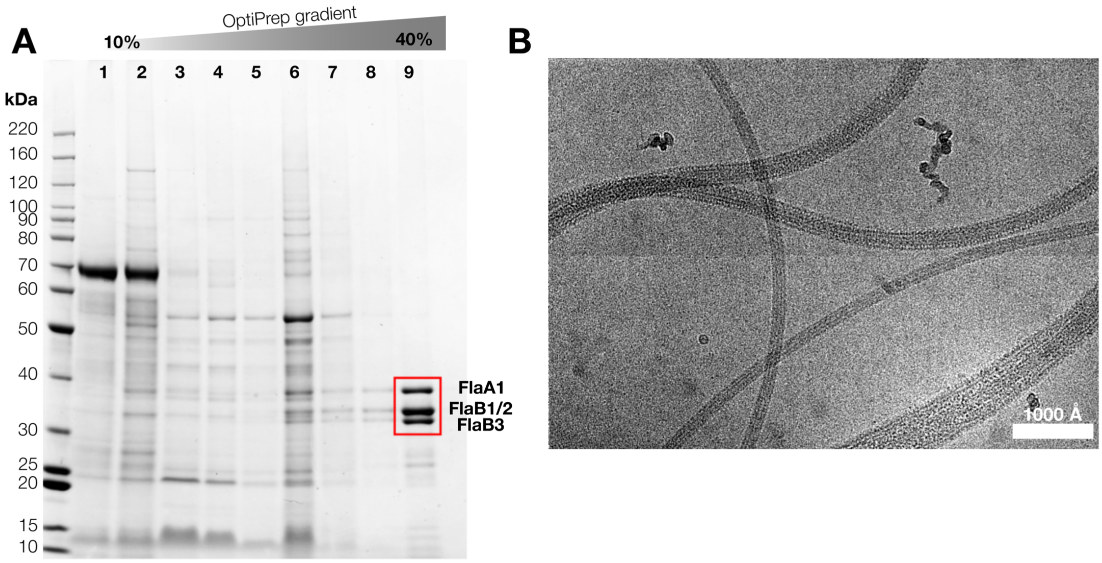


**Supplementary figure 1**. **Purification of *T. denticola* flagellar filaments.** (A) SDS-PAGE analysis of different OptiPrep gradient fractions. Lanes 1–2: 10% OptiPrep fraction; lanes 3–5: 20% fraction; lanes 6–8: 30% fraction; lane 9: 40% fraction containing purified flagellar filaments. The red box highlighted probable FlaA1 (39.3 kDa) and FlaB1/2/3 (31.3/31.6/30.9 kDa) protein bands. (B) Motion-corrected and contrast-enhanced cryo-EM micrograph of the purified flagella.

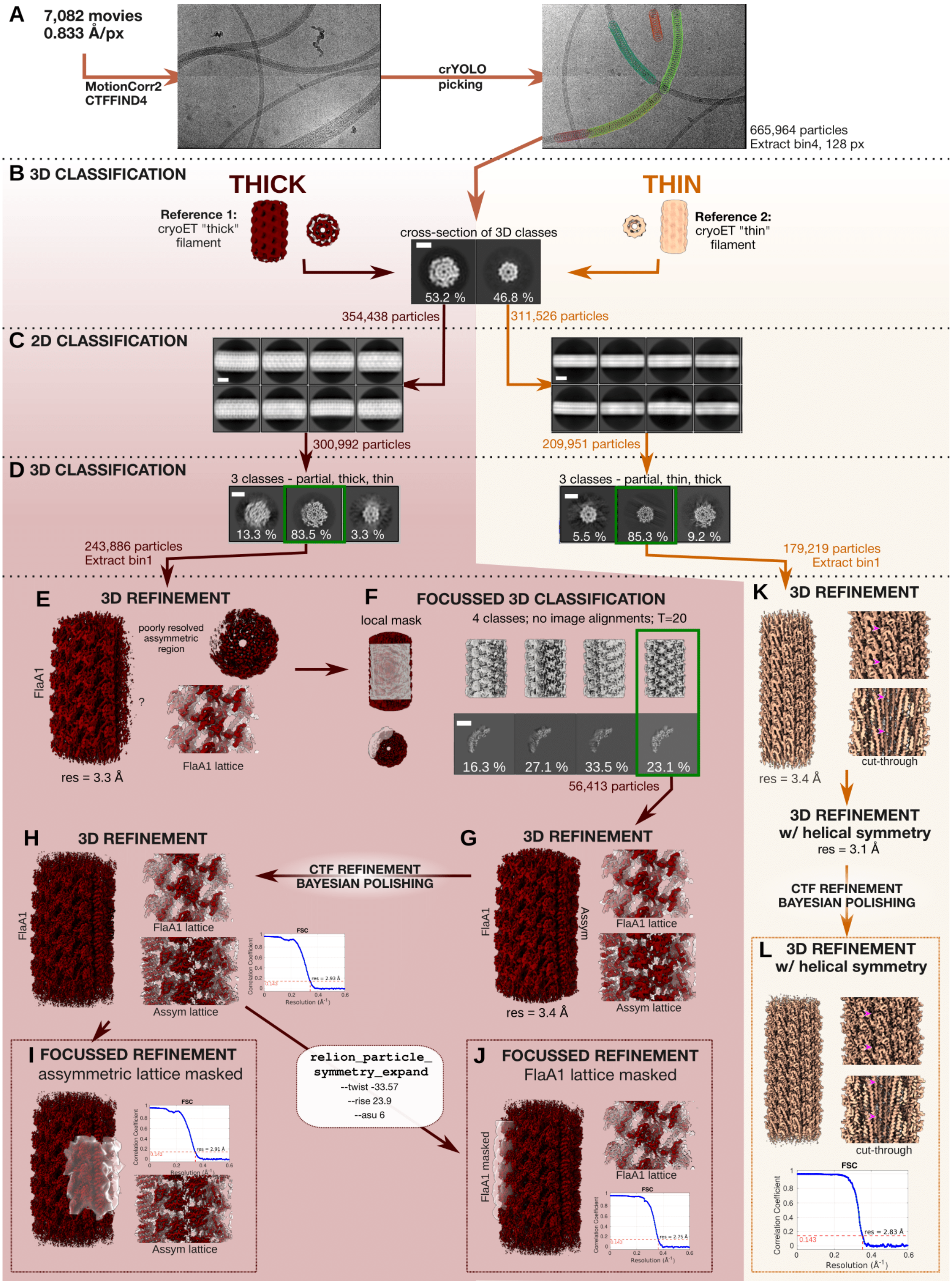
A
7,082 movies
0.833 Å/px
MotionCorr2
CTFFIND4
crYOLO
picking
665,964 particles
Extract bin4, 128 px
B 3D CLASSIFICATION
THICK
THIN
Reference 1:
cryoET "thick"
filament
Reference 2:
cryoET "thin"
filament
cross-section of 3D classes
53.2 %
46.8 %
354,438 particles
311,526 particles
C 2D CLASSIFICATION
300,992 particles
209,951 particles
D 3D CLASSIFICATION
3 classes - partial, thick, thin
13.3 %
83.5 %
3.3 %
3 classes - partial, thin, thick
5.5 %
85.3 %
9.2 %
243,886 particles
Extract bin1
179,219 particles
Extract bin1
E
3D REFINEMENT
FlaA1
poorly resolved
assymmetric
region
?
res = 3.3 Å
FlaA1 lattice
F
FOCUSSED 3D CLASSIFICATION
local mask
4 classes; no image alignments; T=20
16.3 %
27.1 %
33.5 %
23.1 %
56,413 particles
K
3D REFINEMENT
res = 3.4 Å
cut-through
3D REFINEMENT
w/ helical symmetry
res = 3.1 Å
CTF REFINEMENT
BAYESIAN POLISHING
L
3D REFINEMENT
w/ helical symmetry
cut-through
FSC
Correlation Coefficient
res = 2.83 Å
0.143
Resolution (Å⁻¹)
H
3D REFINEMENT
FlaA1
FlaA1 lattice
Assym lattice
CTF REFINEMENT
BAYESIAN POLISHING
G
3D REFINEMENT
FlaA1
Assym
FlaA1 lattice
Assym lattice
res = 3.4 Å
res = 2.93 Å
I FOCUSSED REFINEMENT
assymmetric lattice masked
res = 2.91 Å
Assym lattice
relion_particle_
symmetry_expand
--twist -33.57
--rise 23.9
--asu 6
J FOCUSSED REFINEMENT
FlaA1 lattice masked
FlaA1 masked
FlaA1 lattice
res = 2.75 Å

**Supplementary figure 2. Single particle cryo-EM data processing.** (A) Initial processing and particle picking of >7,000 movies through MotionCorr2, CTFFIND4 and crYOLO allowed extraction of >665,000 particles for processing by RELION-5.0. (B) Initial classification of thick and thin particles through a single iteration of RELION 3D classification with two references derived from the cryo-ET reconstructions (Figure 2). (C-D) The sequestered subsets representative of the two filament types was independently processed through 2D (C) and 3D (D) classification to remove partially assembled or poorly classified particles. (E) Refinement of the thick filaments showed a well resolved inner FlaB core and FlaA1 lattice but a poorly resolved sheath on one side of the filament – labelled the asymmetric region. (F) 3D classification with local masking, a high T threshold and no image alignments separated heterogenous particles into 4 classes. (G) Class 4 (representative of 23% of the particles) was selected for further refinement. (H) After CTF refinement and Bayesian polishing the final resolution for the full lattice was 2.9 Å. (I) Focused refinement of the asymmetric lattice improved local resolution. (J) Symmetry expansion of the FlaA1 lattice followed by focused refinements improved resolution of the FlaA1 sheath to 2.8 Å. (K) For the thin flagella, the first 3D refinement with C1 symmetry resolved to 3.4 Å - pink arrows indicate the helical pitch of individual FlaB within the lattice. Subsequent 3D refinement with application of helical symmetry improved the resolution to 3.1 Å. (L) After CTF refinement and Bayesian polishing the final resolution for the thin filament was 2.8 Å.

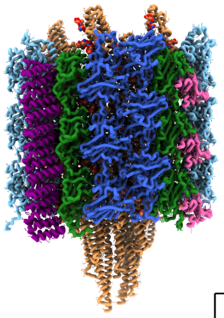

**Assymmetric sheath assembly**
**PDB: 9PIN**
**EMDB: EMD-71673**

**FlaA1 FlaL1 FlaB2**
**FlaA2 FlaL2**
**FlaA3**

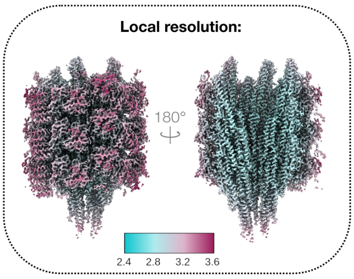


| | Tertiary structure: | Density examples: |
|---|---|---|
| **FlaA1 - TDE1712**<br>**39.3 kDa**<br>**Total AA (incl. SP): 349**<br><br>**Modelled:**<br>**Left - 324**<br>**Right - 318** | 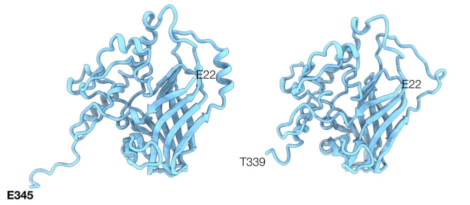  | 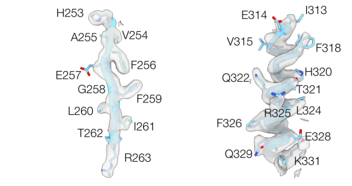  |
| **FlaB2 - TDE1004**<br>**31.6 kDa**<br>**Total AA: 286**<br><br>**Modelled: 286** | 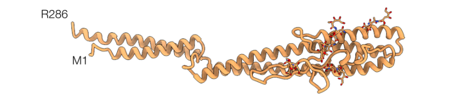  | 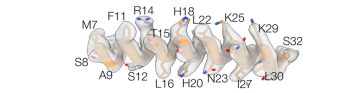  |
| **FlaL1 - TDE2349**<br>**26.2 kDa**<br>**Total AA (incl. SP): 234**<br><br>**Modelled: 190** | 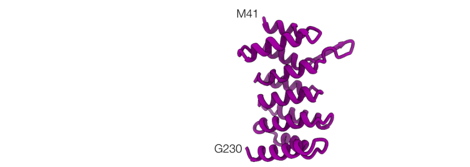  | 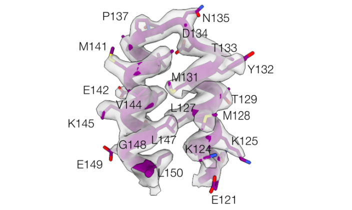  |
| **FlaA2 - TDE1409**<br>**27.8 kDa**<br>**Total AA (incl. SP): 246**<br><br>**Modelled:**<br>**Left - 198**<br>**Right - 221** | 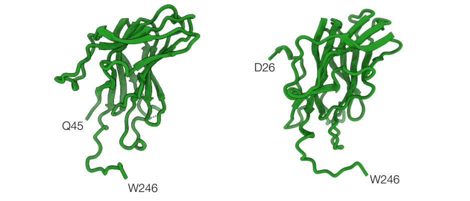  | 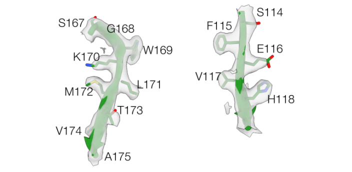  |
| **FlaA3 - TDE1408**<br>**27.1 kDa**<br>**Total AA (incl. SP): 236**<br><br>**Modelled:**<br>**Left - 198**<br>**Right - 200** | 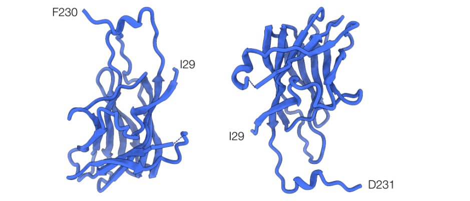  | 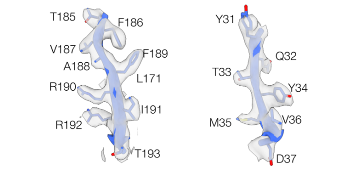  |
| **FlaL2 - TDE1480**<br>**16.1 kDa**<br>**Total AA (incl. SP): 139**<br><br>**Modelled: 110** | 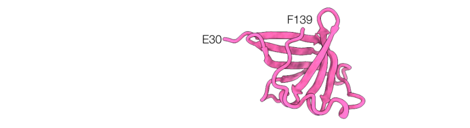  | 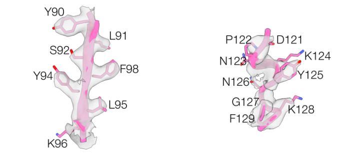  |

**Supplementary figure 3. Model quality for the asymmetric sheath assembly.** The submitted model PDB: 9PIN / EMD-71673 containing 6 copies of FlaA1, 18 FlaB2, 3 FlaL1, 6 FlaA2, 6 FlaA3, and 3 FlaL2. Non-crystallographic symmetry restraints were applied across core flagellins and across each set of three longitudinal repeating components within each model. The laterally repeating copies of FlaA1, FlaA2, FlaA3 were modelled separately. Density reflects the resolution estimates for the reconstruction (2.9 Å) - local resolution estimates (top right) were calculated within RELION-5.0.

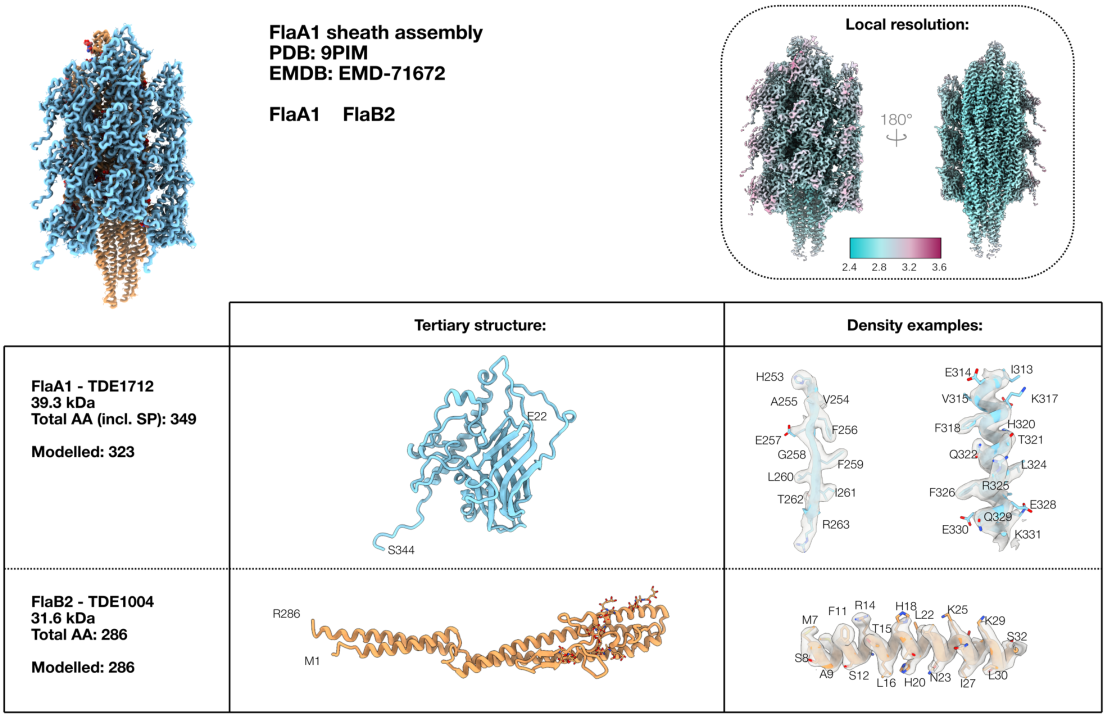


**Supplementary figure 4 – Model quality for the thick FlA1 lattice.** The submitted model PDB: 9PIM / EMD-71672 containing 6 copies of FlaA1 and 9 copies of FlaB2. Non-crystallographic symmetry restraints were applied to all FlaB2 and all FlaA1 thus all repeats are identical. Density reflects the resolution estimates for the reconstruction (2.8 Å) - local resolution estimates (top right) were calculated within RELION-5.0.

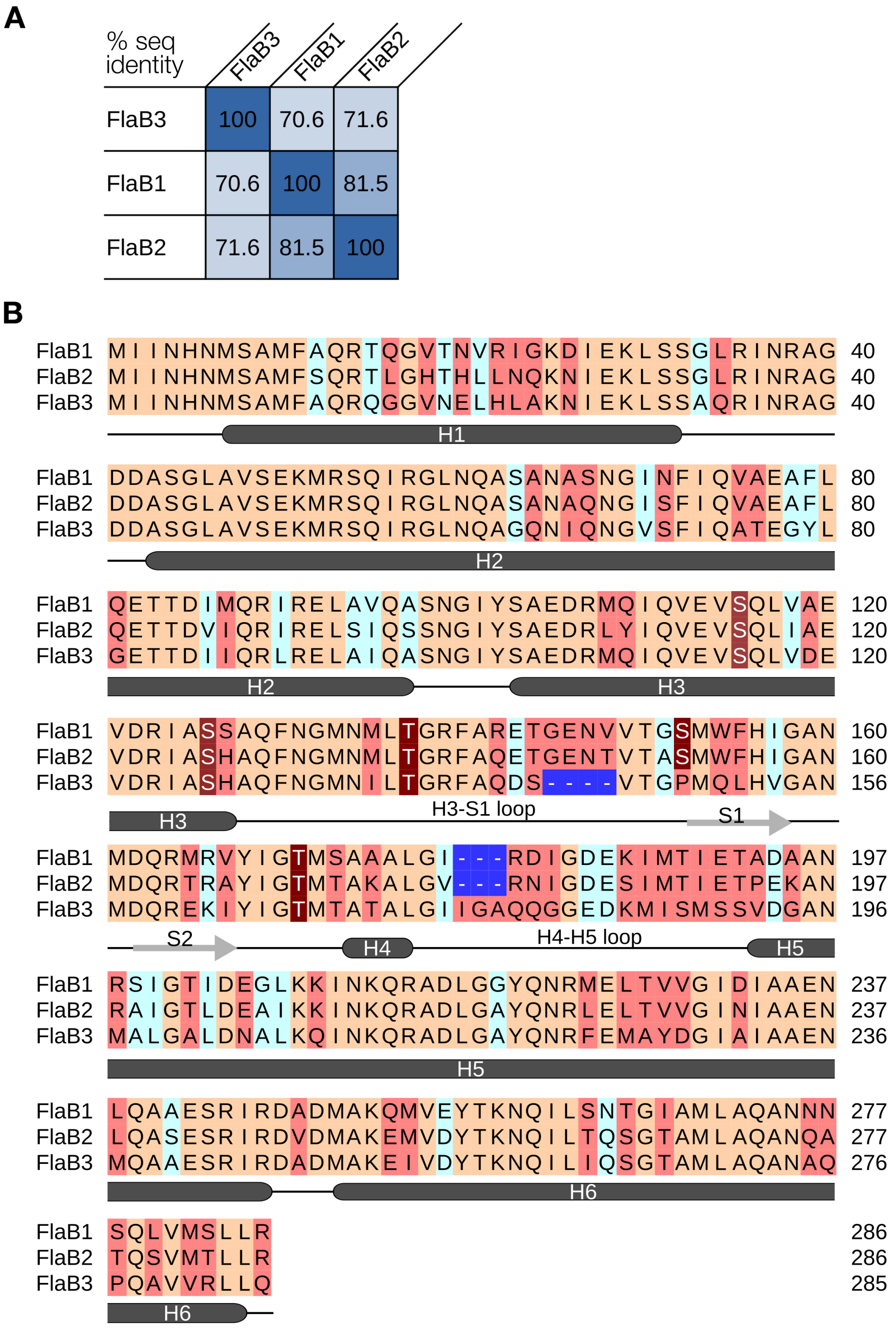


**Supp Fig 5. Homology and sequence analysis for FlaB1, FlaB2, FlaB3 (Uniprot accession #: Q73MN1, Q73NZ6, Q73MN3).** (A) sequence identity matrix for FlaB homologues in *T. denticola*. (B) Sequence alignment of FlaB homologues with secondary features outlined below; α-helices – dark grey; β-stands – light grey.

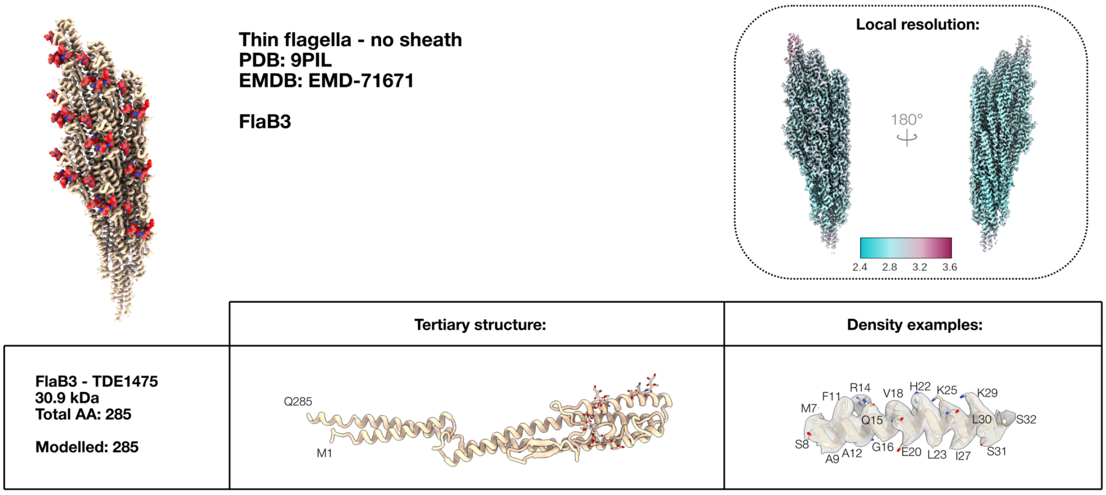


**Supplementary figure 6. Model quality for the FlaB3 thin filament.** The submitted model PDB: 9PIL / EMD-71671 containing 9 FlaB3. Helical symmetry was used during the 3D refinement and non-crystallographic symmetry restraints were applied during model refinement thus the density and models for each FlaB3 should be identical across the lattice. Density reflects the resolution estimates for the reconstruction (2.8 Å) - local resolution estimates (top right) were calculated within RELION-5.0.

### Supplementary tables:

#### Supplementary Table 1 – Mass Spectrometry Data

| Protein IDs | Fasta headers | Mol. weight [kDa] | Intensity | iBAQ |
|---|---|---|---|---|
| TDE1477 | flagellar filament core protein (FlaB1) | 31.31 | 4.78E+09 | 3.37E+08 |
| TDE1712 | flagellar filament outer layer protein (FlaA1) | 39.288 | 5.45E+09 | 3.19E+08 |
| TDE1004 | flagellar filament core protein (FlaB2) | 31.564 | 2.37E+09 | 1.44E+08 |
| TDE1475 | flagellar filament core protein (FlaB3) | 30.945 | 1.23E+09 | 6.42E+07 |
| TDE1409 | flagellar filament outer layer protein (FlaA2) | 27.845 | 8.73E+08 | 5.25E+07 |
| TDE1480 | conserved hypothetical protein (FlaL2) | 16.05 | 7.29E+07 | 9.11E+06 |
| TDE2349 | conserved hypothetical protein (FlaL1) | 26.234 | 8.74E+07 | 8.32E+06 |
| TDE1408 | flagellar filament outer layer protein (FlaA3) | 27.053 | 7.82E+07 | 6.13E+06 |
| TDE2673 | hypothetical protein | 25.504 | 2.58E+07 | 2.87E+06 |
| TDE2508 | hypothetical protein | 50.708 | 5.08E+07 | 2.21E+06 |
| TDE0405 | major outer sheath protein | 58.269 | 6.61E+07 | 1.83E+06 |
| TDE2768 | flagellar hook protein FlgE (flgE) | 49.574 | 3.27E+07 | 1.36E+06 |
| TDE0985 | oligopeptide/dipeptide ABC transporter, periplasmic peptide-binding protein, putative | 75.246 | 4.26E+07 | 8.98E+05 |
| TDE2055 | hemin-binding protein B (hbpB) | 44.806 | 1.70E+07 | 7.71E+05 |
| TDE1717 | hypothetical protein | 23.191 | 6.89E+06 | 6.89E+05 |
| TDE2735 | surface antigen, putative | 49.125 | 8.82E+06 | 4.41E+05 |
| TDE2257 | 5-nucleotidase family protein | 57.836 | 7.79E+06 | 3.71E+05 |
| TDE2056 | outer membrane hemin-binding protein A | 45.185 | 6.49E+06 | 2.82E+05 |
| TDE2352 | flagellar hook-associated protein FlgK (flgK) | 69.535 | 1.93E+07 | 2.60E+05 |
| TDE0626 | hypothetical protein | 52.942 | 3.43E+06 | 1.63E+05 |
| TDE0765 | translation elongation factor Tu (tuf) | 43.788 | 3.75E+06 | 1.56E+05 |
| TDE2601 | surface antigen, putative | 93.848 | 7.09E+06 | 1.54E+05 |
| TDE0842 | cytoplasmic filament protein A (cfpA) | 78.511 | 4.14E+06 | 1.09E+05 |
| TDE1273 | oligopeptide/dipeptide ABC transporter, peptide-binding protein | 60.324 | 3.22E+06 | 1.04E+05 |
| TDE0762 | Serine protease (dentilisin) from NCBI | 77.474 | 6.48E+06 | 8.45E+04 |
| TDE0803 | hypothetical protein | 55.95 | 1.84E+06 | 7.35E+04 |
| TDE0649 | hypothetical protein | 12.667 | 3.23E+05 | 4.61E+04 |
| TDE2119 | glycine reductase complex selenoprotein GrdB2 | 46.302 | 1.17E+06 | 4.32E+04 |
| TDE1341 | conserved hypothetical protein | 52.303 | 7.30E+05 | 2.43E+04 |
| TDE2420 | DNA-directed RNA polymerase, beta subunit, putative | 160.03 | 1.58E+06 | 1.94E+04 |
| TDE1342 | conserved hypothetical protein | 78.573 | 0.00E+00 | 0.00E+00 |
| TDE1422 | glycosyl transferase, group 2 family protein | 40.079 | 0.00E+00 | 0.00E+00 |

**Supplementary Table 2 – CryoEM**

| | **Single particle analysis** | | |
|---|---|---|---|
| magnification | 105, 000 x | | |
| voltage (keV) | 300 | | |
| electron exposure ($e^{-}$/Å$^2$) | 57.7 | | |
| defocus range | -0.5 to –1.8 | | |
| pixel size (Å) | 0.833 | | |
| symmetry imposed | C1 | | |
| movies | 7082 | | |
| particle picking method | crYOLO | | |
| box distance (Å) | 52 | | |
| minimum filament length (particles) | 8 | | |
| threshold | 0.4 | | |
| box size (px) | 448 | | |
| initial particles (total) | 665,964 | | |
| **ARCHITECTURE** | **"THICK"** | | **"THIN"** |
| initial particles (by architecture) | 354,438 | | 311,526 |
| final particles (pre-symX) | 56,413 | | 179,219 |
| **Full map resolution (Å)** | 2.93 | | 2.83 |
| **Masked refinement area** | **Asymmetric lattice** | **FlaA1 lattice** *(after symX w/ 6asu)* | **Full lattice** *(w/ helical symmetry)* |
| **resolution (Å)** | **2.91** | **2.75** | **2.83** |
| FSC threshold | 0.143 | 0.143 | 0.143 |
| **REFINEMENT** | | | |
| sharpening factor | n/a | n/a | n/a |
| map sharpening method | deepEMhancer | deepEMhancer | deepEMhancer |
| model resolution | 2.7 | 2.6 | 2.7 |
| FSC threshold | 0.143 | 0.143 | 0.143 |
| **MODEL COMPOSITION** | | | |
| chains | 63 | 27 | 18 |
| non-hydrogen atoms | 84315 | 44298 | 20475 |
| protein residues | 10425 | 5481 | 2565 |
| ligands (GTP, GDP) | DEN – 90<br>CA - 12 | DEN - 45 | DEN - 36 |
| **B FACTORS (Å$^2$)** | | | |
| protein | 86.80 | 78.02 | 96.54 |
| nucleotide | - | - | - |
| ligand | 133.40 | 118.22 | 146.44 |
| **BONDS (RMSD)** | | | |
| bond length (Å) | 0.002 | 0.001 | 0.001 |
| bond angles (°) | 0.649 | 0.513 | 0.596 |

| | | | |
|---|---|---|---|
| **VALIDATION** | | | |
| MolProbity score | 1.23 | 1.08 | 0.89 |
| Clashscore, all atoms: | 4.54 | 2.87 | 1.48 |
| poor rotamers (%) | 0.45 | 0.79 | 0.45 |
| favoured rotamers (%) | 95.00 | 94.86 | 96.86 |
| **RAMACHANDRAN PLOT** | | | |
| Ramachandran outliers (%) | 0.00 | 0.00 | 0.00 |
| Ramachandran favoured (%) | 98.63 | 98.51 | 98.94 |
| Rama distribution Z-score | 1.75 | 1.83 | 2.97 |
| **DEPOSITION** | | | |
| EMDB | EMD-71673 | EMD-71672 | EMD-71671 |
| PDB | 9PIN | 9PIM | 9PIL |

**<u>Supplementary Information Movie legends:</u>**

**SI Movie 1-** Cryo-EM map of thick flagellar filament displaying different subunits.

**SI Movie 2-** Conformational changes of FlaA1 within the outer sheath of periplasmic flagella.
